# Optimization-Based Synthesis with Directed Cell Migration

**DOI:** 10.1101/2023.12.22.573173

**Authors:** Eric C. Havenhill, Soham Ghosh

## Abstract

Collective behavior of biological agents such as herds of organisms and cells is a fundamental feature in the systems biology and in the emergence of new phenomena in the biological environ-ment. Collective cell migration under a physical or chemical cue is an example of this fundamental phenomenon where individual cell migration is driven by the collective behavior of the neighboring cells and vice versa. The goal of this research is to discover the mathematical rules of collective cell migration using experimental data and testing the predictive nature of the models in independent experimental data. Such insight is made possible in this work with the hybrid use of dynamic mode decomposition (DMD) [1] and optimal control theory. Both single and multi-cellular systems are simulated, including obstacle courses, using this framework. The results of this work show how cells collectively behave during their migration and also, opens the possibility of designing robotic cells for possible therapeutic purpose where the cell trajectory can be controlled.

## I. INTRODUCTION

Collective cell migration is crucial for many biological processes including the formation and regeneration of tissue. Specific examples include osteoblast movement during bone formation, fibroblast movement in wound healing, the movement of macrophages and neutrophils in the immune response and cancer metastasis [2, 3]. Collective migration is also critical in tissue engineering applications where the cells must infiltrate through a scaffold and bypass obstacles to reach an intended target. Collective cell migration can be extended to larger biological systems, such as herd migration in different organisms from bacteria to large animals. Therefore, the development of a predictive biophysical model of the characteristics and key parameters of migrating entities would be beneficial to the scientific community.

This paper outlines a framework for the discovery of some mathematical models of collective cell migration. A governing ordinary differential equation (ODE) for a single cell, or partial differential equation (PDE) for a collection of cells in tandem, is developed and used to model cell migration in the direction defined by a gradient. A hybrid model, based on optimal control, is used in conjunction with the associated ODE or PDE for a synthesis allowing for the study of cell migration in a two-dimensional space.

In literature, DMD has been used to obtain the governing dynamic equations, both ODEs and PDEs, for many well-known systems such as the Van der Pol oscillator and Burgers’ equation [1]. This serves as encouraging evidence that this same methodology will work for biological systems. To illustrate this, consider the diffusion equation in Eq. (1), a parabolic PDE that is common in multiple fields of physics to model phenomena such as thermal, chemical, and particle diffusion. Similar equations, such as the diffusion equation with a different diffusion coefficient and the Fisher-Kolmogorov equation, are being used to model collective cell migration [4–6]. Artificial data can be generated with MATLAB’s pdepe solver of the diffusion equation presented in Eq. (1) with the associated boundary conditions (BC) and initial condition (IC) in Eqs. (2) and (3), respectively.

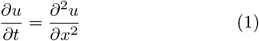

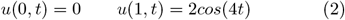

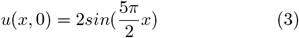

Solving Eqs. (1-3) generates a surface plot shown in Fig. 1. The recovered PDE is also shown for comparison. The successful reconstruction of the governing equations of motion for these dynamic systems provides confidence that the same methodology can be applied to biological systems, specifically for collective cell migration.

**FIG. 1.**
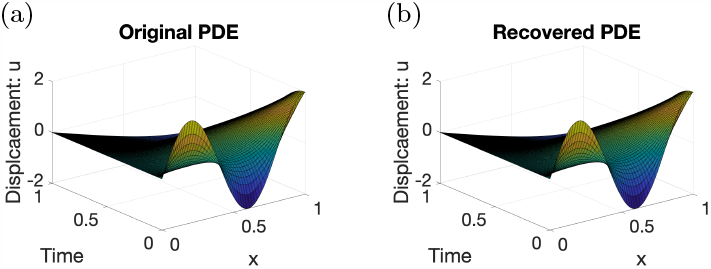
(a) Surface plot generated from Eqs. (1-3). (b) Surface plot generated from the recovered PDE and Eqs. (2-3).

## II. DATA SET

Scratch wound assays were performed by culturing a monolayer of murine fibroblast cell line (NIH 3T3) in an 8-well plate, as described in our recent work [7]. A scratch was applied using a pipette tip. Subsequently, time-lapse image of the cell migration during the wound closure was recorded using a Zeiss LSM 980 confocal microscope with incubation chamber. The automated cell tracking was done on nucblue stained nuclei to collect quantitative data of the cell migration using TrackMate in Fiji[8]. The TrackMate coordinate system and 0^*th*^ data set are illustrated in Fig. 2, where the positive x-direction is horizontal and pointed to the right, and the positive y-direction is vertical and pointed downwards. Methods of directed cell migration include haptotaxis, chemotaxis, topotaxis, and galvanotaxis [9]. From the nature of this experiment, the cells are assumed to migrate due to a chemical attractant, in the direction toward the center of the scratch.

**FIG. 2.**
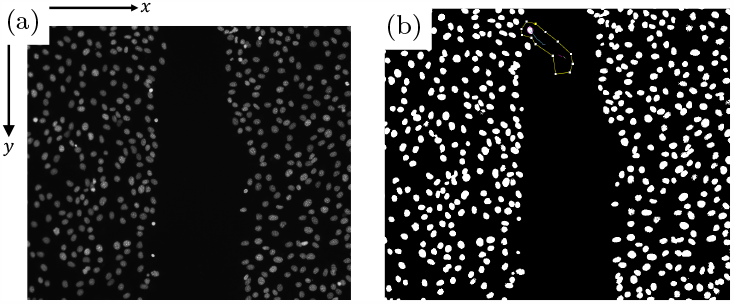
(a) Live cell imaging and automated tracking before wound closure shown of murine fibroblasts for a sample control group illustrating the scratch wound assay. (b) Tracked cell trajectory for an individual cell of interest.

## III. DYNAMIC EQUATIONS

As mentioned, the methodology for discovering the dynamics is outlined in [1] and reproduced in Appendices A, B, C, D, and E for the specific case of the cellular systems here. The larger the library of terms in the A-matrix is, from Eqs. (E2) and (E5), the more robust the model can be, e.g. 1, *O*^1^, *O*^2^, …, *u*′, *u*′′,… Solving this equation will allow the formulation of a differential equation that serves as the governing equation for the cell migration in the x-direction. Thresholding can be done in order to eliminate some of the more insignificant elements of *ξ*. The system solution can be obtained by solving the matrix system shown in Eq. (4).

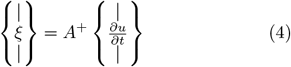

Note that in Eq. (4), *A*^+^ represents the appropriate inverse of choice. The sparse solution for *ξ* was obtained in MATLAB. A sparse or parsimonious solution is preferred since it allows for the identification of the dominant terms that are driving the dynamics. An argument from intuition informs us that the recovered equation should likely contain the fewest number of terms since this is the structure that we see in many important governing PDE’s such as the Korteweg–De Vries equation, diffusion, reaction-diffusion, Burgers, Schrödinger, Navier-Stokes, Kuramoto–Sivashinsky, wave, Laplace, and Poisson equations, to name a few.

### A. Single-Cell System

Now the methodology is ready to be applied to the single-cell and multi-cell systems. The dynamical equations of the single-cell system are shown in Eqs. (F1), (F2), (F4), and (F6). The plots of the trajectory of the cell for the different dynamic systems is shown in Fig. 10.

### B. Multi-Cell System

We seek a method that discovers the partial differential equations of a collective cellular system of the form shown in Eq. (5).

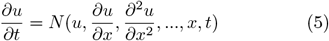

The dynamics are recovered as shown in Eq. (6), where the terms with relatively small coefficients are omitted such that the dominant terms are shown.

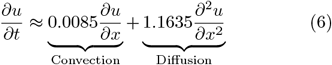

Eq. (6) takes the form of the convection-diffusion equation, however, the sign of the convection coefficient is opposite to the sign of the convection coefficient in the standard convection-diffusion equation. The solution suggests that the convective term serves as a model for some amount of migration of the cells in the direction opposite of the location of the scratch, which is consistent with experimental data; the diffusion term serves as a model for the migration of the cells towards the scratch due to a chemical gradient.

#### 1. a posteriori Dirichlet BCs

The initial condition (IC) for all of the simulations of this PDE is given in Eq. (7).

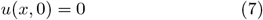

The boundary conditions (BC) are obtained from TrackMate from the trajectories of the cells at the two boundaries. For implementation, the data was interpolated to fit a smooth polynomial with the structure given in Eqs. (8) and (9), where the value of *ν* is chosen according to the user’s discretion.

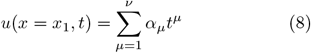

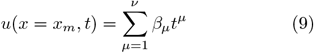

The data of the original experiment is shown in Fig. 3. Comparing the surface plot of the original data with a surface plot of the data generated by the DMD framework shows an overall similar shape.

**FIG. 3.**
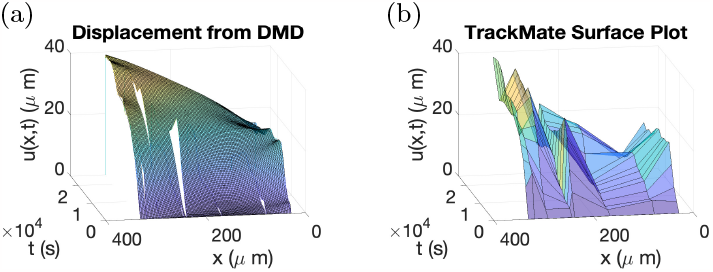
(a) Surface plot of Eq. (6), subject to the BC in Eqs. (8) and (9) and IC in Eq. (7). Waterfall plots of the trajectories obtained from the TrackMate data are included. The final time used in the simulation is the same as of the experiment. (b) Surface plot of the original data. The final time used in the simulation is the same as of the experiment.

#### 2. Velocity-Based Dirichlet BCs

Dirichlet BC can be constructed based on the first few time points of the experiment. The construction of the Dirichlet boundary condition relies upon the assumption that one can obtain the trajectory of the cells at least upon the completion of the first time step, specifically in the form of the time derivative of the position. The first-order time-derivative approximation of the displacement, shown in Eq. (10), can be constructed with a forward-biased stencil, where *δ* represents a difference approximation to a derivative [10].

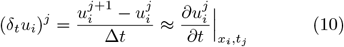

Eq. (10) will be used only when data from the first time step is known, however, for a better approximation to the first derivative, more sophisticated stencils can be used when more data points are accessible from multiple time steps. A couple of examples of these advanced stencils are shown in Eqs. (F7), and (F8). For all of the stencils, the values of *x*_*i*_ are *x*_1_ for the left boundary (wall) and *x*_*m*_ for the right boundary (scratch-wound interface); *j* = 1 in all cases.

In addition to the derivatives, we can develop an approximation of the amount of time that it takes for the cells at the scratch wound interface to reach the center of the scratch, as shown in Eq. (11). For two representative data sets, the final times of the simulation as predicted by Eq. (11) are an order of magnitude larger than the final time of the actual experiment.

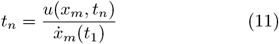

The numerator in Eq. (11) can be experimentally determined once the scratch in the scratch wound assay is produced. Note that for the simulations in Figs. 4 - 6, the final time of the actual experiment was used for the simulation in order to draw a comparison. Using either Eqs. (10), (F7), or (F8), one can develop the boundary conditions that are given in Eq. (12). Note that this same expression is used for the BC at the scratch-wound interface and for the cell at the wall of the well; hence for *x*_*i*_ = *x*_1_ (at the wall) and *x*_*i*_ = *x*_*m*_ (at the scratch-wound interface).

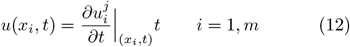

The approach outlined here is anticipated to be the most versatile of all of the approaches, due to the fact that it does not need to know the global position of the cells. The results of this method are shown in Fig. 4.

**FIG. 4.**
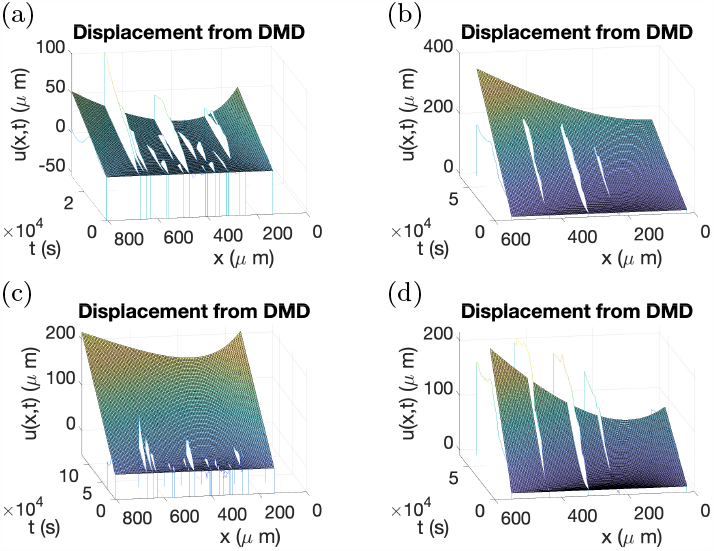
Surface plots of Eq. (6), subject to the BC in Eq. (12) and IC in Eq. (7). Waterfall plots of the trajectories obtained from the TrackMate data are included. For the simulation: (a) The 1^*st*^ data set is used with experiment’s final time. (b) The 1^*st*^ data set is used with experiment’s predicted final time, found from Eq. (11). (c) The 2^*nd*^ data set is used with experiment’s final time. (d) The 2^*nd*^ data set is used with experiment’s predicted final time, found from Eq. (11).

**FIG. 5.**
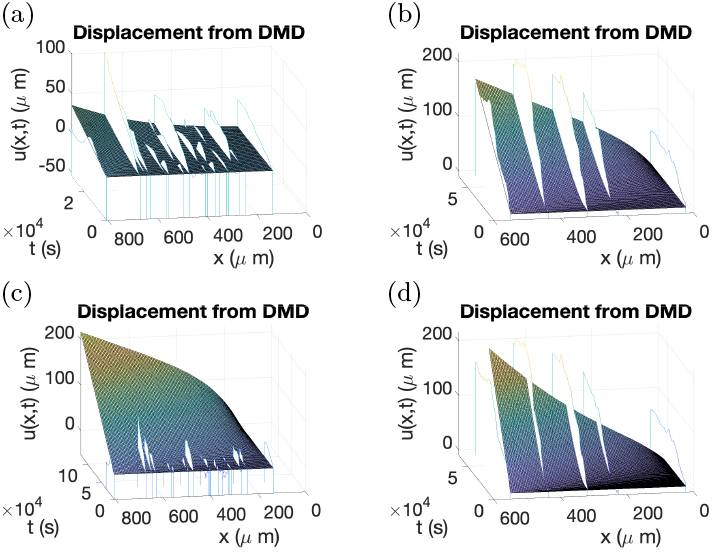
Surface plots of Eq. (6), subject to the BC in Eqs. (12) and (13) and IC in Eq. (7). Waterfall plots of the trajectories obtained from the TrackMate data are included. For the simulation: (a) The 1^*st*^ data set is used with experiment’s final time. (b) The 1^*st*^ data set is used with experiment’s predicted final time, found from Eq. (11). (c) The 2^*nd*^ data set is used with experiment’s final time. (d) The 2^*nd*^ data set is used with experiment’s predicted final time, found from Eq. (11).

**FIG. 6.**
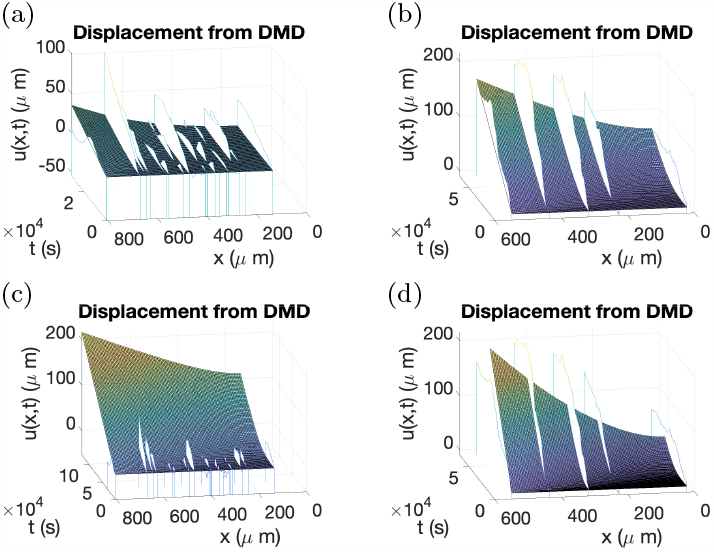
Surface plots of Eq. (6), subject to the BC in Eqs. (12) and (14) and IC in Eq. (7). Waterfall plots of the trajectories obtained from the TrackMate data are included. For the simulation: (a) The 1^*st*^ data set is used with the experiment’s final time. (b) The 1^*st*^ data set is used with the experiment’s predicted final time, found from Eq. (11). (c) The 2^*nd*^ data set is used with the experiment’s final time. (d) The 2^*nd*^ data set is used with the experiment’s predicted final time, found from Eq. (11).

#### 3. Zero-Displacement and Velocity-Based Dirichlet BC

The BC at the right end can be constructed from Eq. (12). The BC at the left end will be assumed to be a zero-displacement BC. The goal here is to provide BCs that will allow the prediction of the cell motion by using the trajectory of the cells during the first time step. The first boundary condition, given in Eq. (13), is obtained by assuming the trajectory of the immediate interior cell to the cell positioned at *x* = *x*_1_ (at the wall).

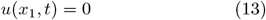

The simulation results of this approach are shown in Fig. 5. It should be noted that for some situations, the BC presented in Eq. (13) can be problematic. For simulations containing a cell whose location with the lowest x-value, *x*_1_, is at the wall of the well, then Eq. (13) is the appropriate BC. However, for cases where the lowest x-value, *x*_1_, is far removed from the wall, then the BC representing zero displacement should not be applied.

#### 4. Mixed (Dirichlet/Neumann) BCs

The BC for the right-hand side from Eq. (12) is used again. The second BC is constructed from the spatial derivative of Eq. (13), as shown in Eq. (14). This BC is generated as an assumption that the neighboring cell to the one at the *x*_1_ location will approximately move the same as the one positioned at the *x*_1_ location.

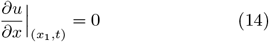

The results of this approach are shown in Fig. 6. This approach is anticipated to serve as the second most robust approach.

## IV. OPTIMIZATION

The movement of the cells in the direction towards the scratch was studied in the previous sections. In order to study the motion of cells perpendicular to the direction of the scratch, which is of particular interest when investigating the 2D trajectory of the cell subjected to obstacles such as other cells, we seek an understanding through an optimization framework. Optimal control relies upon this mathematical framework in order to construct optimal trajectories.

### A. Direct Collocation Method - Optimization Through Dynamic Programming

The optimization problem is formulated in Eq. (15), where the cost functional J to be minimized is the Bolza cost function including the Mayer term, Φ[·], and the Lagrange term, L[·], which is subjected to constraints on the system, where **f** (**x**(*t*), **u**(*t*), *t*), *ϕ*(**x**_1_, *t*_1_, **x**(*t*_*f*_), *t*_*f*_), and **C**(**x**(*t*), **u**(*t*), *t*) represent the dynamics, boundary constraints, and path constraints, respectively. The state vector, **x**, is constructed based on the context of the problem. For example, if the motion of only one cell is of consideration, then **x** ∈ ℝ^2×1^ in order to include both the x and y positions. If the motion of *i* cells is of consideration, then **x** ∈ ℝ^2×1^.

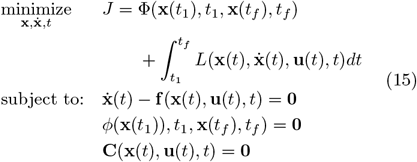

Note that the formulation presented in Eq. (15) is a continuous-time formulation, which means that it needs to be converted into an approximate formulation for the direct collocation through transcription into a nonlinear program. First, the integral in the cost functional to be minimized can be approximated with a quadrature formulation. For example, using the trapezoidal method, the integral given in Eq. (15) can be converted into an algebraic expression. The quadrature conversion used in the simulations presented in this paper are given in Eq. (16).

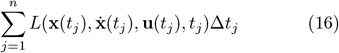

The dynamics will also be discretized, which can be done with a finite differencing scheme such as the one presented in Eq. (D1); however, for better results, MATLAB’s ode45 will be used to numerically integrate the equations. MATLAB’s fmincon function will solve the resulting system of algebraic equations and hence solve the static optimization problems with the use of algorithms such as interior-point, trust-region-reflective, sqp, or others [11]. Finally, once applying this quadrature scheme, we can apply Bellman’s Principle of Optimality, which states that the optimal trajectory between two nodes also lies on the optimal trajectory for the entire trajectory; this will allow us to construct a dynamic optimization problem from many consecutive static optimization problems.

Using this method, several simulations are performed and included in this paper. Simulations without obstacles present are included in Appendix G 2. Simulations with obstacles are presented in IV A 1 and IV A 2.

#### 1. Static Obstacles

For the trajectories generated here, the Lagrangian will be chosen such that the minimum-energy solution is obtained. This is done in order to seek a trajectory for the moving cell that resembles a natural system, since the agents in biological systems tend to expend the least amount of energy. With the initial position of the cell and the displacement data obtained from solving the PDE in Eq. (6) subject to the IC and BC of choice, DMD can be used in order to generate an ODE that describes the motion of the cell’s x-directional movement. An example ODE resulting from this process is shown in Eq. (17).

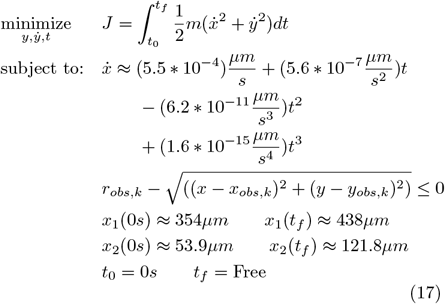

The optimization can be performed in the presence of obstacles, specifically static cells. This simulation of this is shown in Fig. 7.

**FIG. 7.**
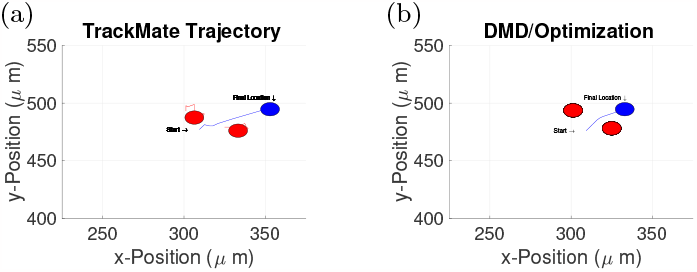
(a) Actual trajectory of the cell of consideration, as obtained from TrackMate. (b) The hybrid DMD-optimization model, resulting in an optimal trajectory, with the discovered dynamics from Eq. (17).

#### 2. Dynamic Obstacles

The simulations in Fig. 7 assume that the obstacles are static, and hence the obstacles don’t move through the entirety of the simulation. Using the results from the previous sections, a more realistic simulation is included for comparison to the static case. The other cells are assumed to move according to the PDE discovered from the DMD. The controlled cells move according to an optimization policy, specifically a minimum energy approach. Each cell is allowed to arrive at the goal at their own final time; they do not have to arrive at the goal location at the same time. The results of different dynamic optimizations are shown in Fig. 8. Similar simulations have been done [12] with the use of a potential field approach, however, such approaches do not use an optimization-based framework.

**FIG. 8.**
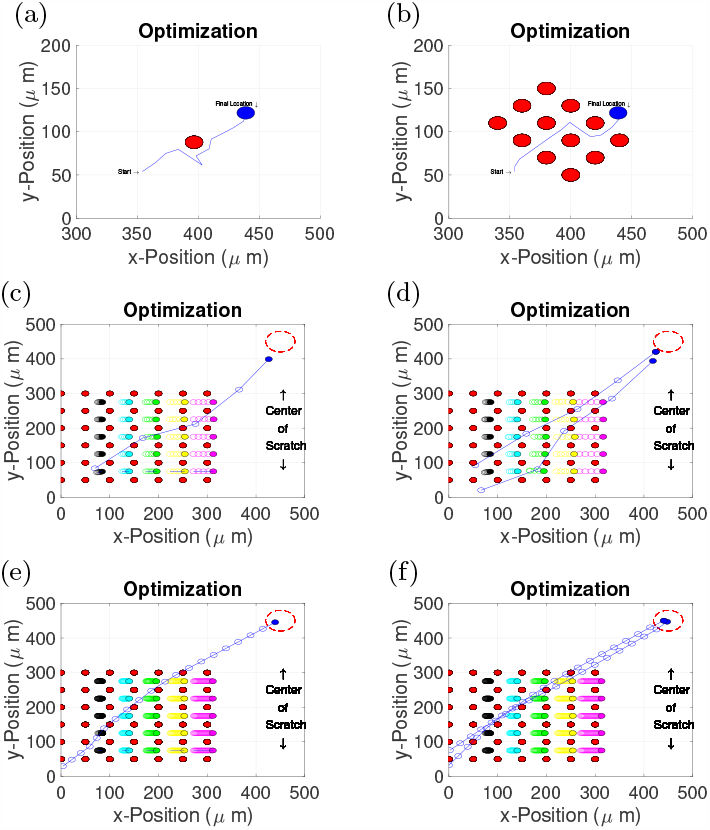
(a) An individual cell’s traversing to a prescribed final location. (b) An individual cell’s traversing through an obstacle course of static obstacles. (c), (d), (e) and (f) An individual or multiple cell or cells traversing through an obstacle course of both moving and non-moving obstacles.

## V. DISCUSSION

The outcome of this paper demonstrates a new framework for the discovery of the rules of cell migration through the synthesis of a hybrid approach with DMD and optimal control. The mathematical model frames the cell migration in terms of a predictive displacement field, rather than on population density, which is the approach practiced by most current models. Like many PDEs in physics, the discovered PDE only relies upon a few terms to reasonably model the dynamics. While many terms were allowed to be in the dynamics, only a few terms appear in the result.

The DMD was executed in this work, which allows for the successful description of the one-dimensional movement of a single cell and multiple cells. This is different than a linear interpolation of the migrational data, since it relies upon existing spatio-temporal data and hence cannot predict the global level population behavior. While the governing equation shown in Eq. (6) contains a diffusion equation that represents the migration of the cells towards the scratch, a convection term is also present that represents bulk migration in the direction opposite to the scratch, possibly attributed to the bulk retraction of the cell monolayer during the migration. When simulated, the PDE will have the same IC as presented in Eq. (7), however the BC will vary depending on the context of the experiment; Eq. (12) can be used when the global position of the cell furthest from the scratch is unknown, Eqs. (12) and (13) can be used when the cell furthest from the scratch is known to be located at the well wall; Eq. (13) and (14) can be used when the other methods fail due to limitations in calculating the time derivatives at the beginning of the experiment. However, it should be noted that there are several limitations and challenges to the application of DMD that were experienced in this paper. While the DMD can be used with a noisy system, here represented by small Brownian motion of a cell during migration, the recovered governing equation will not predict such random fluctuations. Another limitation of DMD in the context of this experiment relates to the availability of a fine spatio-temporal grid on which the finite differencing is performed. These limitation arise from the number of cells (relevant for spatial differentiation) and the number of time points due to the allowable sampling frequency of a cell system with a microscope (relevant for temporal differentiation). Due to these constraints on the system, the solution of the PDE shown in Fig. 3 is imperfect, whereas the recovered system in Fig. 1 much more closely resemble the original system. It should also be noted that for cell migrational systems, the construction of a governing equation with this DMD process is not always possible, particularly due to the availability of only a coarse spatio-temporal grid.

Using a hybrid DMD/optimal control-based model for a single cell with nearby obstacles, an optimal path in two-dimensions was developed. The simulated cell in Fig. 7 does not move as far in the x-direction as the cell from the experiment due to the large displacement of this particular cell in the experiment; the actual cell’s movement is abnormally large. A minimum-energy path is still generated with a notably similar trajectory to the experimental cell’s trajectory.

The optimization was further performed to predict the optimal path in a dynamic environment in two dimensions for one cell or multiple cells. This optimization shows the capability of predicting the optimal path of a cell in two-dimensions with respect to the other cells in the population that are following the path defined by the DMD. Observing the control policy generated by the optimization, a bang-bang control type is obtained for the cells that are controlled.

The initialization for the states of the optimization is very important for obtaining meaningful results for methods like those presented in this paper. The final solution of the optimization can be sensitive to the initial guess; with a poor guess, the solution can converge to an inferior minimum. For this paper, the cells for which the optimal trajectory was generated were given an initial guess for their trajectory which is formed from a linear interpolation of their initial position to their goal location. Since the cells are modeled as first-order systems here, this is a reasonable assumption. For refinement, h-methods were used to attenuate the error and thereby converge the solution to the optimal trajectory; p-methods were not considered due to the capabilities of numerically integrating with MATLAB’s ode45.

The hybrid DMD/optimization method is limited due to a high computational cost. Since the time to complete the simulation on a 2018 MacBook Pro with 16 GB of RAM is approximately one hour and 24 hours for the simulations presented in Fig. 8 (e) and (f), respectively, this method cannot be used quickly in real-time in the current state. However, with a more powerful computer it can be possible. Since the computational cost rises significantly with the number of nodes, one possible way to alleviate this long simulation time is to perform a simulation with a fewer number of nodes and then use a method that is not computationally expensive, such as gradient descent with potential functions, in order to guarantee an obstacle avoidance.

A simulation of a second-order moving cell was not included due to needing the quantification of the force bounds with which the cells exert themselves on the substrate. Data from this experiment provided velocity data only; however, a future analysis can help to quantify these cell traction forces and the system may be resimulated for refinement.

Practical implementations of such a model exist in the field of medical treatment services that involve magnetic micro-robots, such as drug delivery with a micro-robot [13]. Research has been done to develop automated control and path planning of micro-robots [14]; it is expected that the research outlined in this paper will allow for the solving of a similar problem but from a different approach. The optimization framework presented in this paper will also open possibilities into research related to studying cell migration at the individual level.

## ACKNOWLEDGMENTS

The authors wish to acknowledge the collection of the scratch-wound assay data by Briar Hine and Jack Forman.

## Appendix A: DMD

In order to illustrate these concepts here, a popular dynamic system of consideration from the field o f physics, the Van der Pol Oscillator, will be considered in Eq. (A1), where 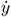, *ÿ*, and *μ* represent the time derivatives of the state, second time derivative of the state, and coefficient, respectively.

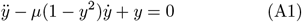

A synthetic data set of this system can be generated in MATLAB, allowing the production of spatiotemporal data ofthe two states of the system, *x*_1_ = *y* and 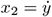. Using the methodology already mentioned in the literature, as will be reproduced in this paper, the ODE can be recovered [15]. The A-matrix in Eq. A2 is used, which contains a library of functions that are assumed to participate in the system’s dynamics.

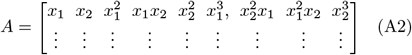

The plot of the original dynamics compared to the recovered dynamics are shown in Fig. 9.

**FIG. 9.**
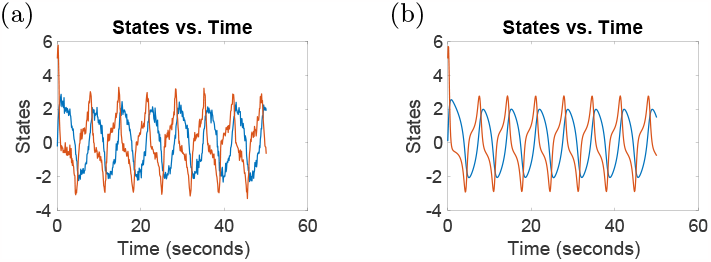
(a) Plot of the states of the Van der Pol oscillator with noise added to the system. (b) Recovered states of the Van der Pol oscillator recovered dynamics from the DMD process.

## Appendix B: Data Set

### 1. Single-Cell System

The data obtained from TrackMate is arranged in an arbitrary order, where n represents the number of time points. An example of this is shown in Eq. (B1).

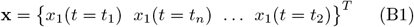

### 2. Multi-Cell System

The original data is gathered in a matrix similar to the following, where the columns are arranged in an arbitrary order. The *i*^*th*^ column represents the x-values of the trajectory of the *i*^*th*^ biological cell in the system. The *j*^*th*^ row of a *i*^*th*^ column represents the *i*^*th*^ biological cell’s x-value at the *j*^*th*^ time point. An example of this is shown in Eq. (B2).

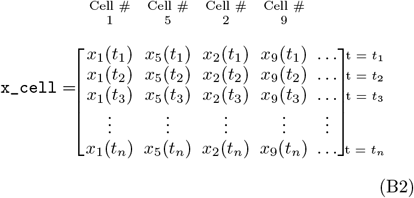

For the multi-cellular case, m represents the number of cells we are using and n represents the number of time points that we have for each cell. In general, the positions are monotonically increasing as in Eq. (B3), and for all strictly increasing time as in Eq. (B4).

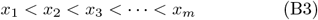

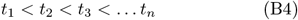

## Appendix C: Dynamic Model Discovery - Sol’n. Formulation

### 1. Single-Cell System

#### a. Organize the Data

The data of the positional points over time will be organized into the vector that is shown in Eq. (C1). The subscript on the x-value represents the biological cell’s number. The same process for the y-direction will be repeated for this cell, but not shown here to avoid redundancy.

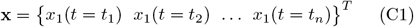

### 2. Multi-Cell System

#### a. Organize the Data

The matrix of positional vs. temporal data for each cell is given in Eq. (C2).

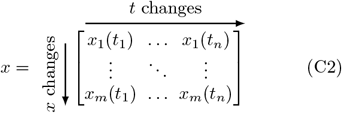

The matrix of displacement data is given in Eq. (C3).

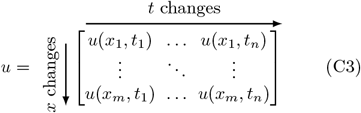

This is the displacement matrix that corresponds to the displacement field of the cells, where the displacement is defined as the difference between the current x-position of the biological cell and the original x-position of the cell, as shown in Eq. (C4), where *i* = 1, 2, …, *m* and *j* = 1, 2, …, *n*.

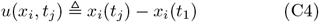

## Appendix D: Finite Difference Method

Both spatial and temporal derivatives of the data will need to be calculated since they will be going into the matrix in Eq. (E5). Finite differencing is used in order to calculate the derivatives, however, it should be noted that more sophisticated differentiation techniques, such as total variation of derivatives, can be used in order to improve the model. For this paper, simple differencing schemes, such as Euler backward and the central difference method, as opposed to higher-order stencils, are used to calculate the first and second-order derivatives, respectively; although they are simple stencils, they are sufficient to perform the differentiation.

### 1. Single-Cell System

The finite differencing is outlined in the multi-cell system solution, so the process is not repeated here in order to avoid redundancy.

### 2. Multi-Cell System

#### a. First-Order Differentiation - Time Derivation

The finite difference differentiation scheme for a first-order time derivative is given in Eq. (D1).

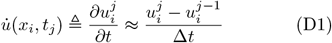

For implementation in this paper, the differentiation stencil expressed in Eq. (D1) will be expressed equivalently in matrix form in Eq. (D2).

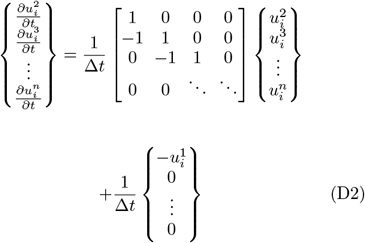

The time derivatives are calculated for each biological cell throughout its entire trajectory, which will result in a differencing matrix, shown in Eq. (D2). After each cell’s trajectory is numerically differentiated with respect to time, they are assembled into the matrix given in Eq. (D3), where 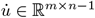.

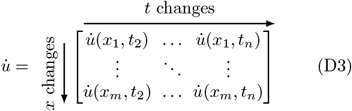

However, since we will be using higher-order derivatives in time, we will also omit the last column from this matrix, so that the dimensions remain compatible. The first and last rows will also be omitted, due to the same reason, but for the spatial derivatives. Therefore, we will get an entire matrix for the time-derivative of *u*, 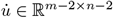, that takes the form given in Eq. (D4).

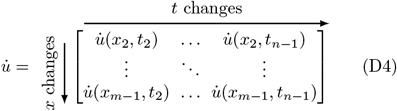

#### b. First-Order Differentiation - Spatial Derivation

The finite difference differentiation scheme using Euler backward for a first-order spatial derivative, which assumes that the initial condition is known, is given in Eq. (D5).

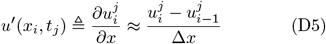

For implementation in this paper, the differentiation stencil expressed in Eq. (D5) will be expressed equivalently in matrix form in Eq. (D6).

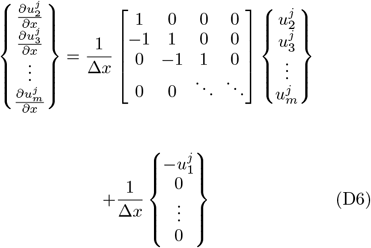

#### c. Second-order differentiation - Spatial Derivation

The finite difference differentiation scheme for the central difference method for a second-order derivative is given in Eq. (D7).

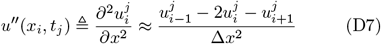

For implementation in this paper, the differentiation stencil expressed in Eq. (D7) will be expressed equivalently in matrix form in Eq. (D8).

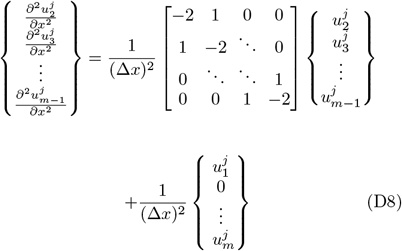

## Appendix E: DMD Formulation

### 1. Single-Cell System

The finite differencing techniques for the singular cell will be similar to those done for the multi-cellular system. Both systems use a finite differencing scheme to approximate the derivative at particular nodes, which will be needed in Eq. (E2).

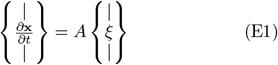

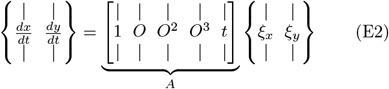

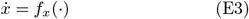

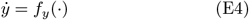

A form of matrix A was originally shown in Eq. (A2) for the Van der Pol system; here, the matrix is chosen at the discretion of the engineer. Sample matrices are given in Section III. The system is solved, as shown in Eq. (4). The following list will illustrate the appropriate choice of inverse when solving this system of equations.

1. If A has fewer rows than columns, then the system has infinite exact solutions that can be found using the pseudo-inverse.
2. If A is square and full-rank, then the standard inverse can be used.
3. If A has more rows than columns, then no exact solution exists, but an approximate one can be found using the pseudo-inverse.

### 2. Multi-Cell System

With time-history data of the trajectories and the derivative data of the cells obtained from TrackMate, illustrated in Fig. 2, and the differentiation from the finite difference method, the multi-cellular system of equations can be assembled, shown in Eq. (E5).

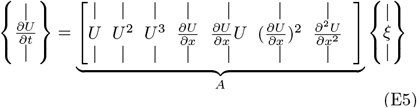

The structure given in Eq. (E5) is made possible by reshaping the matrices into vectors that are stacked on top each other, as is done in Eq. (E6), for example, where *u* ∈ ℝ^*m*−2×*n*−2^ and *U* ∈ ℝ^(*m*−2)∗(*n*−2)×1^.

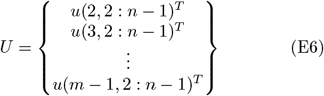

## Appendix F: Dynamic Equations

### 1. Single-Cell System

Several different structures for the A-matrix are explored in this Subsection in order to study their effects of the accuracy of the recovered dynamics. The data from the trajectory illustrated in Fig. 2 is used for this study.

#### a. Structure of A - Version 1

Using a matrix with one term, shown in Eq. (F1), the dynamics are recovered as shown in Eq. (F1).

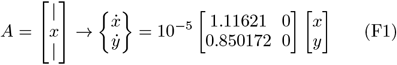

#### b. Structure of A - Version 2

Using a matrix with two terms, shown in Eq. (F2), the dynamics are recovered as shown in Eq. (F2).

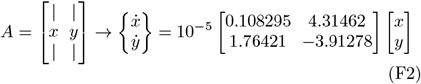

#### c. Structure of A - Version 3

The matrix containing four different terms is shown in Eq. (F3).

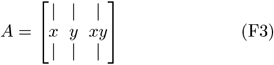

The dynamics are recovered as shown in Eq. (F4).

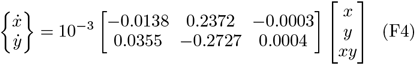

#### d. Structure of A - Version 4

The matrix containing four different terms is shown in Eq. (F5).

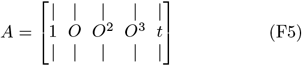

The dynamics are recovered as shown in Eq. (F6).

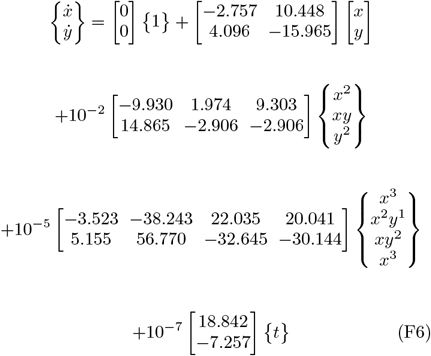

The plots of the original TrackMate trajectory data, in addition to the plots from the solution of the dynamics subject to the boundary conditions in Table I obtained from the DMD, are shown in Fig. (10), where for the graphs on the right-hand side, the blue and red curves represent the x and y-positions of the cell, respectively. It’s apparent that the trajectory from the DMD-based dynamics approaches the trajectory of the cell obtained by TrackMate as the number of approximating terms in the dynamics increases. This gives further confidence that the PDE regression solution will give reliable results.

**TABLE I.**
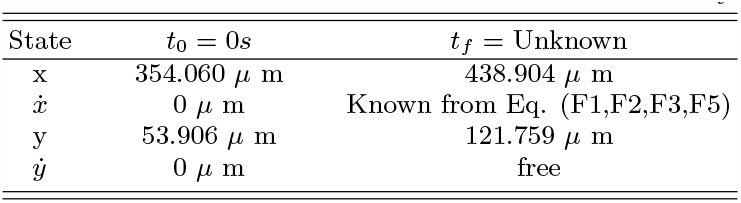
The values of the states and time at the boundary.

### 1. Multi-Cell System

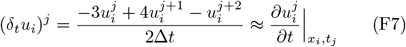

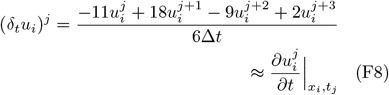

## Appendix G: Optimization

### 1. ODE Optimization

The path constraint, **C**(**x**(*t*), *t*) is assumed to be zero, since there will be no obstacles. The kinetic energy of the cell is defined in Eq. (G1), where *m*_*i*_ represents the mass of the *i*^*th*^ cell, *I*_*i*_ represents the inertia tensor of the *i*^*th*^ cell, 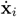 represents the velocity of the *i*^*th*^ cell, and *ω*_*i*_ represents the angular velocity of the *i*^*th*^ cell.

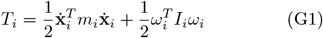

The relationship between force and the potential is defined in Eq. (G2), where *F*_*i*_ represents the traction force of the *i*^*th*^ cell, *V*_*i*_ represents the potential field of the *i*^*th*^ cell, and *u*_*i*_ represents the displacement of the *i*^*th*^ cell as defined in Eq. (C4).

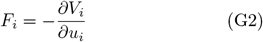

Values of the traction forces during cell migration range from 0 nN to 120 nN [16]. The traction force will be assumed to linearly decrease from this maximum to the minimum value during its migration. The potential field is modeled in Eq. (G3), where the *c*_1_, *c*_2_, and *c*_3_ terms arise as constants of integration and when considering just one cell, *i* = 1.

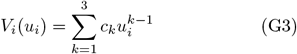

For the case of the cell’s motion that is governed by the ODE, Table I gives the values that are used in the simulation. The choice of 1, T, and T-V for the L(·) function is to seek to find a minimum time, energy, and action, respectively, for the cell migrational problem. The mass of the system is given as m=2.29 ng [17]. The velocity bounds in Eq. (G4) are obtained from experimentation.

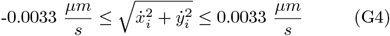

The optimized single-cell system is shown in Fig. 11. The x and y directional motion of the cells are fixed subject to the discovered dynamics; the vertical motion is free. The endpoints are fixed in time and space. The results of these simulations from the hybrid synthesis presented serve as good predictors of cell movement that are at the interface of the scratch. However, it is still desirable to be able to predict the movement of the cells that are located on the scratch and wall boundaries.

**FIG. 10.**
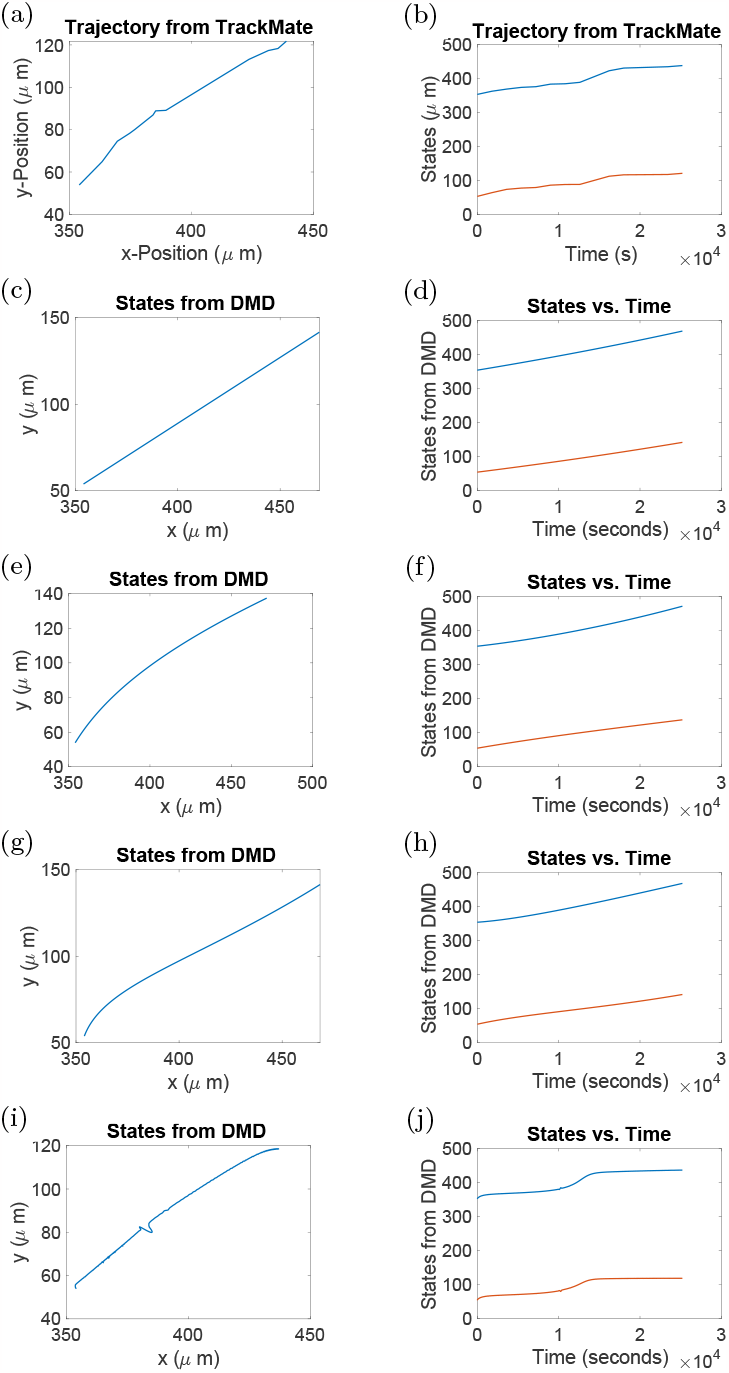
Plots showing the increase in accuracy of the recovered dynamics for comparison. For the plots on the right-hand side, the blue and red curves represent the x and y-positions of the cell, respectively

**FIG. 11.**
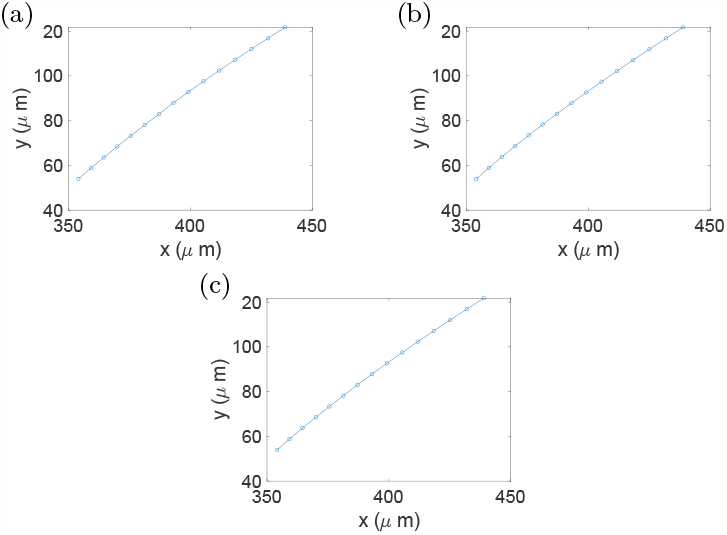
Graphs of the hybrid DMD-optimization trajectories for the single cell, with a Lagrangian cost of (a) time. (b) energy. (c) action.

### 2. Optimization and Prediction of Multiple Cell Trajectories

With the framework developed, the trajectories of multiple cells can obtained. The obstacles could be represented by other cells or fibers in the extracellular matrix.

## Notes

### Competing Interest Statement

The authors have declared no competing interest.

